# Speed Control and Force-Vectoring of Blue Bottle Flies in a Magnetically-Levitated Flight Mill

**DOI:** 10.1101/351445

**Authors:** Shih-Jung Hsu, Neel Thakur, Bo Cheng

**Affiliations:** Department of Mechanical and Nuclear Engineering, Pennsylvania State University, University Park, PA 16802, USA

**Keywords:** insect flight, flapping flight, Calliphora, helicopter model

## Abstract

Flies fly at a broad range of speeds and produce sophisticated aerial maneuvers with precisely controlled wing movements. Remarkably, only subtle changes in wing motion are used by flies to produce aerial maneuvers, resulting in little directional tilt of aerodynamic force vector relative to the body. Therefore, it is often considered that flies fly according to a helicopter model and control speed mainly via force-vectoring enabled primarily by body-pitch change. Here we examine the speed control of blue bottle flies using a magnetically-levitated (MAGLEV) flight mill, as they fly at different body pitch and with different augmented aerodynamic damping. We identify wing kinematic contributors to the changes of estimated aerodynamic force through testing two force-vectoring models. Results show that in addition to body pitch, flies also use a collection of wing kinematic variables to control both force magnitude and direction, the roles of which are analogous to those of throttle, collective and cyclic pitch of helicopters. Our results also suggest that the MAGLEV flight mill system can be potentially used to study the roles of visual and mechanosensory feedback in insect flight control.

Summary statement: This paper elucidates how files control flight speed while flying in a magnetically-levitated (MAGLEV) flight mill, which enables the manipulation of body pitch and aerodynamic load.

## INTRODUCTION

Flies are eminent miniature flyers that exercise stable and agile flight over a large flight envelop (Beatus et al., 2015; Fry et al., 2003; Muijres et al., 2014). This aerial success hinges partly on flies’ ability to precisely control subtle wing movement through regulating the firing rate and timings of steering muscles (Dickinson and Tu, 1997; Lindsay et al., 2017), despite that their wings are difficult locomotor apparatus to control neuromuscularly (Balint, 2004; Deora et al., 2015) or to emulate in engineering designs (Keennon et al., 2012; Ma et al., 2013; Roll et al., 2013). Previous research has shown that even subtle changes of wing motion are sufficient to produce large maneuvering moment for a fly to execute rapid maneuvers, for examples during saccade (Fry et al., 2003), evasive maneuvers (Muijres et al., 2014) and recoveries from aerial stumbles (Ristroph et al., 2010). However, such subtle changes only result in little directional changes of aerodynamic force vector relative to the body; therefore, flies maneuver mostly according to the a helicopter model (Medici and Fry, 2012; Muijres et al., 2014), although they are able to produce large modulation of wing motion through the clutch and gearing mechanisms at the wing hinge (Deora et al., 2015).

The helicopter model may also apply to forward flight, as the flight speed of flies and other insects is well known to tightly correlate with its body pitch angle (David, 1978; Dudley and Ellington, 1990; Meng and Sun, 2016; Willmott and Ellington, 1997), suggesting that the tilt of aerodynamic force vector might also be small during forward flight. However, key questions remain: What wing kinematic variables do they use to control flight speed? How do these variables vary with body pitch and thrust force? and to what degree do flies change the magnitude and direction of aerodynamic forces while flying at different speeds? While the answers to these questions remain elusive, they are of critical importance for insect flight research and also for inspiring novel engineered flight, especially considering that flies and other insects fly at a broad range of speeds and produce large linear acceleration during foraging, chasing mates and escaping from predators (Collett and Land, 1975; Dudley, 2000). For example, locusts *Nomadacris septemfasciata* reach a speed of 13 m/s in a wind tunnel (Waloff, 1972), drone-fly *Eristalis tenax* 8.5 m/s (Meng and Sun, 2016), hawkmoth *Manduca sexta* 5 m/s (Willmott and Ellington, 1997) and bumblebee *Bombus terrestris* 4.5 m/s (Dudley and Ellington, 1990). Dragonfly *Plathemis lydia* accelerates near 2g in prey interception flights (Mischiati et al., 2014) and blue bottle flies *Calliphora vicina* demonstrate 3g acceleration in a free flight chamber (Bomphrey et al., 2009).

Forward flight of insects is commonly studied in laboratory settings using wind tunnels and flight mills. Using wind tunnels, past studies range from the observation of body and wing kinematics in free (Azuma and Watanabe, 1988; David, 1978; Dudley and Ellington, 1990; Meng and Sun, 2016; Willmott and Ellington, 1997) and tethered flight settings (Vogel, 1966), identifying visual control principles (Baird et al., 2005; Fry et al., 2009; Medici and Fry, 2012; Srinivasan et al., 1996), and kinematic-data-driven modeling of flight control and stabilization (Fuller et al., 2014). Flight mills - devices that approximate continuous forward flight in a confined space through restricting an insect to a circular flight path around a pivot joint - are commonly used to determine the traveling distance of insects and their dispersal potentials (Attisano et al., 2015; Ranius, 2006; Ribak et al., 2017). Although flight mills are rarely used to study other aspects of insect forward flight, they have the potential to provide more naturalistic visual and proprioceptive sensory feedback than wind tunnel experiments. This is important because vision plays a key role in regulating forward flight speed. Previous studies have found that many insects (e.g., honeybees and flies) can robustly extract their ground speed (or retinal slip velocity) from visual patterns of varying spatial and temporal frequencies (David, 1982; Fry et al., 2009). Flies are also shown in the wind tunnel experiments to sometimes maintain a preferred ground speed invariant to substantial changes in airspeed (David, 1982) (putatively detected by air flow sensors, e.g., antenna (Fuller et al., 2014)). Therefore, flies, possibly other insects also, are able to fly at their preferred speed independent of aerodynamic power requirement if it is within their locomotor capacities. However, it is unknown whether such behavior can be reproduced in the flight mill experiments, how insects control their flight speed in the flight mill, or what happens when the limit of their locomotor capacity is reached.

In this study, we examined the speed control of blue bottle flies (*Calliphora vomitoria*, N=5, 42.0 ± 8.9 mg) in forward flight using a novel magnetically-levitated (MAGLEV) flight mill. The MAGLEV flight mill, which eliminated the mechanical friction of pivots and permitted systematic manipulation of a fly’s body pitch angle and aerodynamic damping, enabled us to study the details of speed control and force-vectoring and the corresponding wing kinematic control in forward flight. In particular, we tested two force-vectoring models and determined the wing kinematic contributors to the changes in the magnitude and direction of aerodynamic forces.

## MATERIALS AND METHODS

### MAGLEV flight mill apparatus

The MAGLEV flight mill apparatus was comprised of four main components (Fig. 1A): 1) three magnetically-levitated permanent magnets as a pivot joint; 2) a horizontally-rotating shaft with attached fly and damper; 3) inner and outer enclosing walls with grating patterns and 4) three highspeed video cameras (Fastcam Mini UX100, Photron, Japan). The magnetic levitation was achieved through two electromagnets as actuators that stabilized the vertical position of the permanent magnets (i.e., the pivot joint) and rotating shaft using positional feedback provided by two linear Hall-effect sensors (A1321, Allegro microsystem, LLC. Worcester, MA, USA). The first Hall-effect sensor was placed slightly above the permanent magnets to measure the total magnetic field of the permanent magnets and electromagnets combined. The second Hall-effect sensor was attached to the rim of the top electromagnet to separate the noise (magnetic field of electromagnets) from the first Hall-effect sensor. The strength of the magnetic field was then transformed to distance as a proximity signal. A proportional-integral-derivative (PID) controller computed the current compensations for the two electromagnets to keep pivot pin/rotating shaft vertically stable. All sensor readings and computations were processed with a microcontroller (Uno, Arduino, Italy).

**Figure 1.**
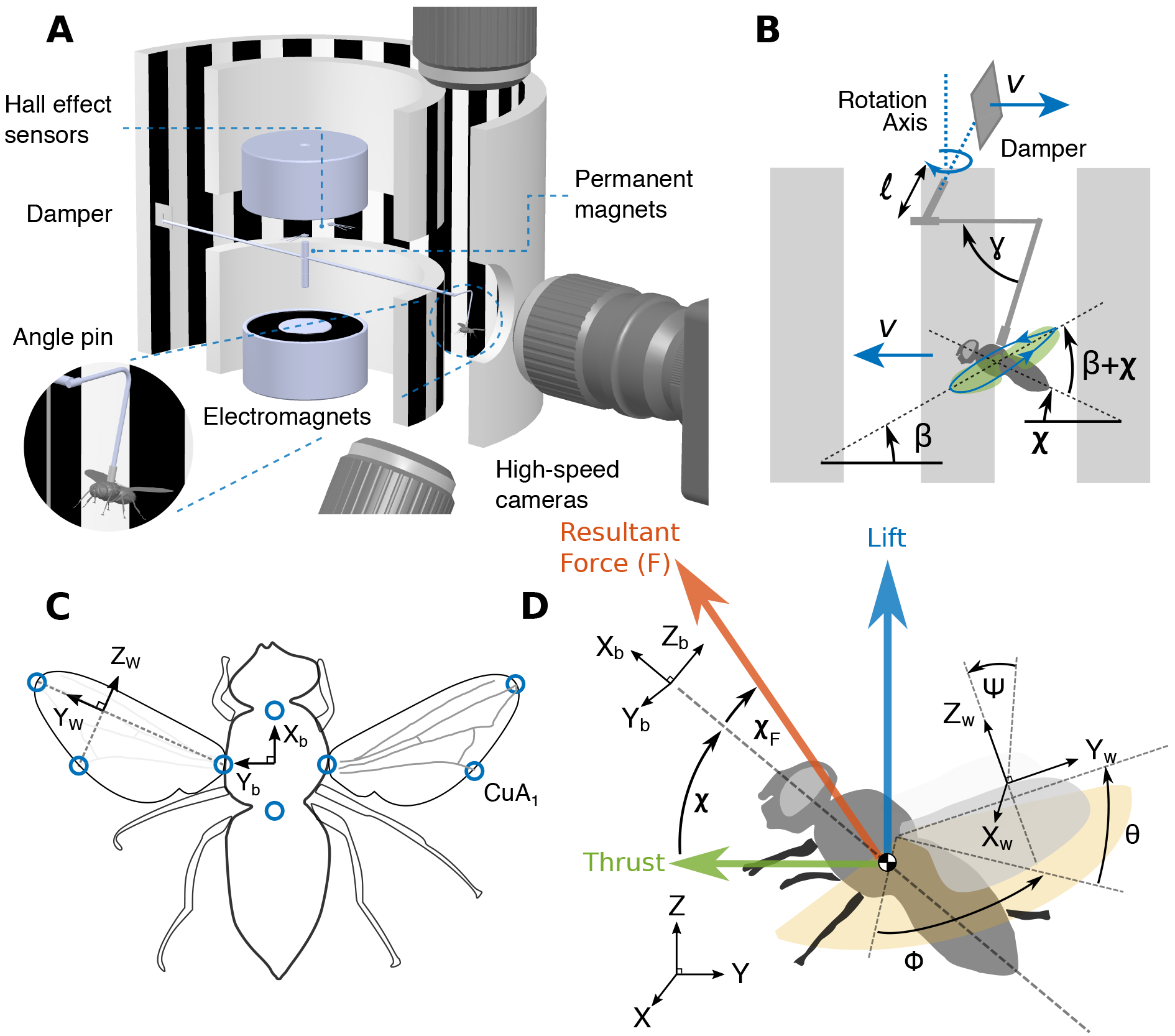
Experimental apparatus, coordinate frames and kinematic variables. (A) Apparatus of the MAGLEV flight mill. MAGLEV flight mill is comprised of electromagnets, permanent magnets, a shaft (carbon fiber, 2 × 254mm, diameter × length), Hall effect sensors, aerodynamic dampers, angle pins and a microcontroller (Uno, Arduino). The entire setup is surrounded by enclosed walls with grating patterns to provide a consistent visual reference. Three high-speed cameras are used to capture body and wings movements of blue bottle flies in forward flight. (B) The angle pins give rise to angle γ between the horizontal plane and a fly’s normal body axis. *χ* is the actual body pitch angle (*χ* ≈ 90° −γ) measured from DLTdv6 (Hedrick, 2008). *β* is the angle between the stroke plane and the horizontal plane. *l* is the radius of the carbon fiber shaft. (C) Anatomical landmarks (blue circles) used for body and wing kinematic extraction. (D) Definitionsof Body frame (*X*_*b*_, *Y*_*b*_, *Z*_*b*_), wing frame (*X*_W_,*Y*_W_,*Z*_*W*_), stroke plane (yellow shade area), wing stroke (*ϕ*), wing rotation (*ψ*), and wing deviation (*θ*). Cycle-averaged lift is represented by the blue arrow parallel and opposite to the direction of gravity (−*Z*). Cycle-averaged thrust is represented by the green arrow orthogonal to the lift vector in and lies in *X*_*b*_ − *Z*_*b*_ plane. The resultant force (red arrow) is the sum of the lift and thrust vectors.

The shaft was made of a 2 × 254 mm (diameter × length) carbon fiber rod, which was sandwiched between the permanent magnets. On one end of the shaft, a magnetic metal angle pin connected a blue bottle fly to the shaft through two micro permanent magnets (1.58 × 3.18 mm, diameter × length) (Fig. 1A-B) glued to the fly’s dorsal. Caution was taken in the gluing process to minimize interference to a fly’s thoracic movements and to maintain a constant angle between the angle pin and the fly’s body. Note that exact body pitch angle (*χ*) was calculated from DLTdv6 (Hedrick, 2008) instead of angle pin angle (γ) since subtle difference existed among flies and each angle pin placement (Fig. 1B). With permanent magnet attached on dorsal, different magnetic angle pins can be easily switched to provide different prescribed pitch-angle. On the other end of the shaft, a damper was attached to create additional drag that the fly needed to overcome in steady forward flight. Two dampers of different sizes (D1: 12.4 × 12.7 × 2.5 mm and D2: 15.2 × 16.2 × 2.5 mm, width × length × thickness), together with the no damper case (D0, where the aerodynamic drag only came from the shaft and the insect body), total three augmented aerodynamic damping conditions were used in the experiments.

Flies mainly rely on visual feedback to regulate their flight speed. To provide a consistent visual environment and to enhance the visual cues they receive, an inner cylinder wall (diameter 203.2 mm) and an outer cylinder wall (diameter 304.8 mm) with identical square wave grating patterns (50.8 mm interval) were used to enclose the flight mill. As a result, the flies flew in the circular corridor (width 50.8 mm) between the two walls (Fig. 1A).

To record the body and wing movements of the flies, three synchronized high-speed cameras were placed on the top, bottom and sideways of an enclosure region spanning approximately 50 × 50 × 50 mm of the circular flight corridor (Fig. 1A). A circular hole was cut on the outer wall for the sideways camera to see through the corridor. We illuminated the enclosure region with three 100W LED light (MonoBright LED Bi-color 750, Genaray, Brooklyn, NY, USA). The video resolution was set to be 1280 × 1024 pixels with 4000 s^−1^ frame rate and 8000 s^−1^ shutter rate. Cameras were calibrated using direct linear transformation for three-dimensional body and wing kinematics extraction (Hedrick, 2008).

### Animal preparation

We used 4- to 7-day-old blue bottle flies (*Calliphora vomitoria*) hatched from pupae purchased commercially (Mantisplace, Olmsted Falls, OH, USA) and cultured in the laboratory. For each experiment, we first cold anesthetized the flies in a refrigerator for 10 minutes (Duistermars and Frye, 2008) and then transferred them to a tethering stage on an oval notch plate with dorsal side up. We used UV cure glue (4305, Loctite Corp.) to attach the micro-permanent magnet on the dorsal side of the thorax. Next, the flies were put to rest to recover from anesthesia for one hour. We then attached the flies to the rotating shaft of the flight mill and started the experiments.

### Experimental procedure

We first tested the flight performance of the flies on the flight mill and only those that could complete at least five laps of flight with 45° angle pin and the largest damper (D2) were used for the experiments. Each fly was attached to one end of the rotating shaft with angle pins held at 0°, 22.5°, and 45° (Fig. 1B). The actual body pitch angles measured from the experiments were 5.5° ± 3.9°, 25.2° ± 3.3° and 41.2° ± 6.2°. The slight differences between the angle pin angle and the body pitch angle were mainly due to the slight misalignment of the angle pins to the normal of the flies’ thorax. For each angle pin, three aerodynamic damping conditions described above were tested. To initiate the flight, a gentle puff of wind gust was introduced to the fly. After the initiation of flight, a fly reached to a constant forward flight speed when the wing thrust was balanced by the total aerodynamic drag acting on the damper, shaft and fly’s body. After at least five laps of flight, we started recording using high-speed cameras (sample recordings are available in supplementary materials S3). In total, all flies had to complete nine different conditions (three angle pins and three damping conditions). For each condition, at least four repeated trials were performed, and for each trial at least four wingbeat cycles were recorded. After completing the trials within one condition, the flies were removed from the flight mill, put to rest for at least 10 minutes, and fed with sugar water before being used for the next condition.

### Damping calibration

The damping coefficients of the combined damper, rotating shaft and a fly’s body for nine different conditions were calibrated using free responses of the rotating shaft. For each calibration, a dead fly with its wings removed was attached to the shaft using one of the three angle pins. To initiate the free response of the rotating shaft, a wind gust was applied to the damper. We recorded timing profile (*t*) from the start to stop with a microcontroller of the rotating shaft triggering two photodiodes spaced with known distance and calculated its angular velocity (ω). Using blade-element analysis (Leishman, 2006) to model the aerodynamic drag, which is assumed to be quadratic, it can be shown that the equation of motion of the shaft is:

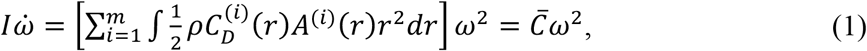

where *I* is the moment of inertia of the shaft, damper and the fly’s body, 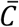 is the calibrated damping coefficient, *ρ* is air density, *m* is the total number of objects that contribute to drag force (e.g., the shaft, dampers, and insect body), *i* is the index of an object. For *i*^*th*^ object, *A*^*i*^(*r*) is the cross-sectional area of a blade-element at radial distance *r* from the shaft center of rotation and 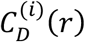 is the corresponding drag coefficient. Integrating Eqn. 1 yields the theoretical speed profile of the shaft:

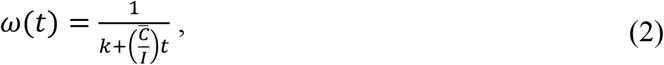

where *k* is the constant of integration. We then performed least square curve fitting to obtain 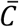. The mean, standard deviation, and coefficient of determination *R*^2^ are reported in Table 1.

**Table 1.** Results of damping coefficient calibration. N, M and L represent no damper, medium and large damper cases, respectively.

| Body pitch angle $\chi$ (°) | 0 | 22.5 | 45 | 0 | 22.5 | 45 | 0 | 22.5 | 45 |
| --- | --- | --- | --- | --- | --- | --- | --- | --- | --- |
| Damper | N | N | N | M | M | M | L | L | L |
| $\bar{C}$<br>($10^{-7} \cdot \text{Kg} \cdot \text{m}^2/\text{rad}$) | 3.79<br>$\pm 0.41$ | 3.89<br>$\pm 0.36$ | 3.81<br>$\pm 0.23$ | 6.25<br>$\pm 0.42$ | 5.44<br>$\pm 0.26$ | 5.98<br>$\pm 0.43$ | 8.18<br>$\pm 0.77$ | 7.53<br>$\pm 0.38$ | 7.45<br>$\pm 0.68$ |
| $R^2$ | 0.99 | 0.98 | 0.99 | 0.99 | 0.99 | 0.98 | 0.99 | 0.99 | 0.99 |

### Kinematics extraction

We used DLTdv6 (Hedrick, 2008) to digitize anatomical landmarks on the body and wings of the blue bottle flies (Fig. 1C), which were then used to calculate the body velocity, pitch angle and wing angles (Fig. 1C-D). We defined the body roll axis (*X*_*b*_) as a unit vector from Thorax-abdomen junction to Head-thorax junction, pitch axis (*Y*_*b*_) from left wing base to the right wing base, and yaw axis (*Z*_*b*_) using the cross-product of *X_b_* and *Y*_*b*_. We defined wing spanwise axis (*Y*_*w*_) as a vector from wing base to wing tip. The cross-product of *Y*_*w*_ axis with a vector from wing base to the trailing edge location of vein (CuA_1_) determined the wing normal *X*_*w*_ axis. Next, the crossproduct of *X*_*w*_ and *Y_w_* determined wing chordwise axis *Z*_*w*_. Body and wing rotation matrices were then calculated based on the corresponding body and wing principal axes (Murray et al., 1994), respectively. A stroke plane frame (*X*_*S*_, *Y*_*S*_, *Z*_*S*_) was defined by rotating the body frame about *Y*_*b*_ axis where *X*_*S*_ intersected with the maximum and minimum sweep positions formed by the wing base-wing tip vectors.

Wing Euler angles (stroke position (*ϕ*), stroke deviation (*θ*) and wing rotation (*ψ*), Fig. 1D) were calculated from the wing rotation matrices (Murray et al., 1994) (from the stroke plane frame to the wing frame) and body pitch angles were calculated from the body rotation matrices (Fig. 1D). Body translational velocities about each principal axis were calculated by taking the derivatives of the head-thorax junction positional vector. Time series of body and wing kinematics were calculated for four complete wingbeat cycles. For each trial, we then calculated time-averaged wing kinematics from the four wingbeat cycles, while also averaging the left and right wing kinematics (mirrored with respect to *X*_*b*_ − *Z*_*b*_ plane). Euler angles of each trial were parameterized using a fifth-order Fourier series prior to kinematics analysis:

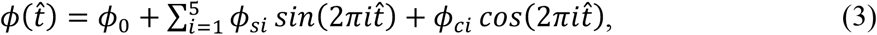

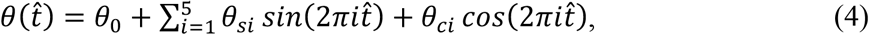

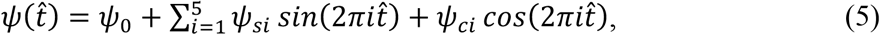

where 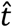 is the dimensionless time of a wingbeat cycle (ranging from 0 to 1); *ϕ*_0_, *θ*_0_ and *ψ*_0_ are constant terms and *ϕ*_*si*_, *ϕ*_*ci*_, *θ*_*si*_, *θ*_*ci*_, *ψ*_*si*_ and *ψ*_*ci*_ are Fourier sine and cosine coefficients and *i* is the order of the Fourier series.

As flies change their continuous wingbeat trajectories to modulate aerodynamic forces and moments, the key changes can be captured by a finite number of wing kinematic variables that represent certain cycle-averaged features (Faruque and Sean Humbert, 2010; Sun, 2014; Taylor, 2001). Here we selected 9 distinct variables that were potentially involved in the speed control and tested their contribution in the force-vectoring models (next section). The 9 wing kinematic variables were: 1) mean wingbeat frequency (*n*), 2) ratio of downstroke and upstroke durations (*T*_*d*_/*T*_*u*_), 3) stroke amplitude (Φ), 4) rotation amplitude (Ψ), 5) deviation amplitude (Θ), 6) mean stroke angle 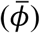, 7) mean rotation angle 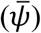, 8) mean deviation angle 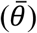, and 9) stroke plane angle (*β*; + *χ*).

### Constant and variable force-vectoring models and variable importance

Here we developed two force-vectoring models for the speed control of flies flying steadily in the MAGLEV flight mill: 1) constant force-vectoring model and 2) variable force-vectoring model. During steady flight, the torque acting on the shaft of the flight mill was zero, which meant that the torque due to the thrust created by the flapping wings (*τ*_*T*_) was equal to those due to the aerodynamic drag of the shaft, damper and insect body combined (*τ*_*D*_), the latter was proportional to the linear speed of the flies or the angular velocity of the shaft. Assuming the flapping wings create a cycle-averaged aerodynamic force with a magnitude of *F* and an angle *χ*_*F*_ from the body longitudinal axis (Fig. 1D), it can be shown that

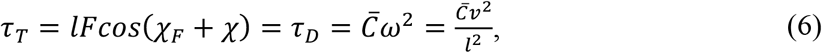

where *l* is the radius of the pivot of the flight mill shaft, and *ν* is the linear velocity of the fly. The constant force-vectoring model can be derived by assuming both *F* and *χ*_*F*_ were constants, i.e., *F*_0_ and *χ*_0_, respectively; therefore the forward velocity can be predicted by,

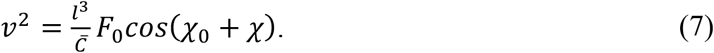

To estimate the values of *F*_0_ and *χ*_0_ from the body kinematic data, a nonlinear least-square regression model was used, where the residual sum of squares *(RSS)* is minimized:

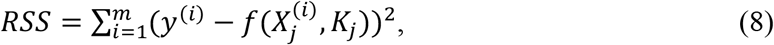

where *m* is the number of trials, 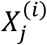 (*i* = 1~*m* and *j* = 1~2) is a vector of known variables including damping coefficients 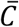 and body pitch angle *χ* from *i*^*th*^ trial, *K*_*j*_ (*j* = 1~2) represents a vector of regression coefficients to be estimated, i.e., *F*_0_ and *χ*_0_, and *y*^(*i*)^ is the square of forward velocity of *i*^*th*^ trial. We performed the nonlinear regression using MATLAB Statistics and Machine Learning Toolbox (Matlab, The MathWorks, Inc., Natick, MA, USA) to estimate parameters *K*_*j*_.

In the variable force-vectoring model, it was assumed that both *F* and *χ*_*F*_ also depended on a collection of wing kinematic variables, e.g., stroke amplitude, frequency, mean rotation angle. Therefore, it was assumed that 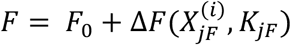 and 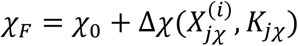, where 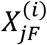 and 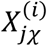 are vectors of wing kinematic variables, the values of which were known from the wing kinematic data; and *K*_*jF*_ and *K*_*jχ*_ are the regression coefficients for the wing kinematic variables, and *F*_0_ and *χ*_0_ are the regression constant terms. Δ*F* and Δ*χ* are the changes of force magnitude and direction due to wing kinematic variables, which were assumed as linear functions. Accordingly, Eqn. 8 can be rewritten as:

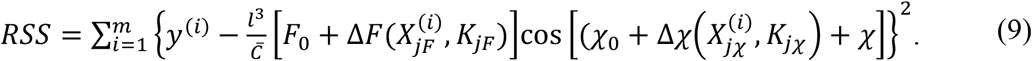

Note that in this regression process, we standardized each variable by subtracting its mean and then dividing by its standard deviation. This standardization rendered all variables on the same metric so that the regression coefficients were not influenced by the variables’ standard deviations (O’Rourke et al., 2005). We assumed that the changes of force magnitude (Δ*F*) depend on 8 kinematic variables (out of the 9 variables mentioned above), i.e., 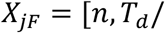 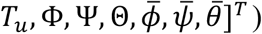; and the changes of force direction (Δ*χ*) also depend on 8 kinematic variables, i.e., 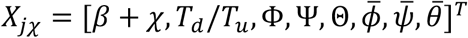. Note that 7 out of the 9 variables are shared between Δ*F* and Δ*χ*, while the stroke plane angle (*β* + *χ*) exclusively affects the force direction and wingbeat frequency (*n*) exclusively affects the force magnitude. As a result, a total of 16 variables were included in the variable force-vectoring model.

The complexity of the variable force-vectoring model depended on the number of wing kinematic variables used. It is well-known that model with overly large number of parameters suffers from overfitting that could overinterpret the data (Burnham and Anderson, 2003). Therefore, model selection using Akaike information criterion (*AIC*) (Akaike, 1998) was performed to evaluate the trade-off between the goodness-of-fit and model complexity. *AIC* is defined as:

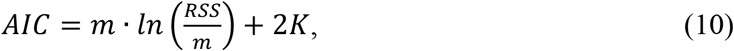

where *K* is the number of parameters in a candidate model. As a rule of thumb (Burnham and Anderson, 2003), the small-sample-size corrected version of Akaike information criterion (*AIC*_*c*_) is preferred if 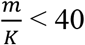. *AIC*_*c*_ is defined as:

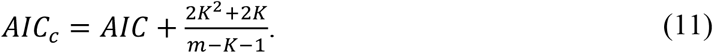

Then, we calculated the *AIC*_*c*_ difference (Δ_*i*_) between the 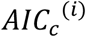 of the *i*^*th*^ candidate model and the minimum *AIC*_*c*_ of all models 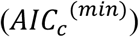,

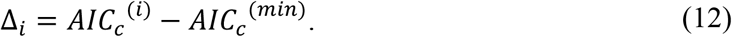

The relative likelihood of *i*^*th*^ candidate model (*g*_*i*_), given the wing kinematic data *X*_*jF*_ and *X*_*jχ*_ for *i*^*th*^ model can be computed as,

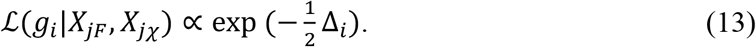

Next, Akaike weights (*w*) for all model combinations were calculated to quantify the importance of each wing kinematic variable. The Akaike weight (*w*) of *i*^*th*^ candidate model is defined as:

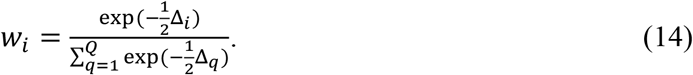

We then summed the Akaike weights over the subset of models that included *X*_*j*_ variable and ranked the variable importance based on the summations of Akaike weights (*w*_+_). In addition to *AIC*_*c*_, Bayesian information criterion (*BIC*) was also calculated for evaluating the trade-off between the goodness-of-fit and model complexity (Burnham and Anderson, 2003), which is defined as:

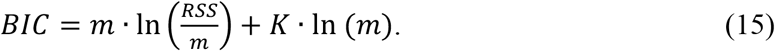

Note that with the natural logarithm of the trial number (*m*), *BIC* applies a larger penalty compared to *AIC*_*c*_ to the model complexity when *m* increases, which tends to result in simpler models.

With calibrated damping coefficients 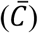, forward velocity (*ν*), and the best-approximating model, cycle-averaged thrust (*F*_*Thrust*_) and lift (*F*_*Lift*_) can be estimated according to,

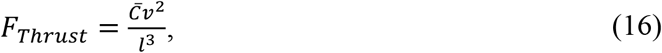

and

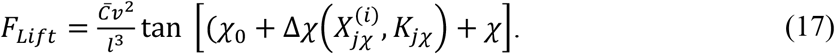

## RESULTS

### Forward flight speed and its dependency on body pitch angle and aerodynamic damping

Using three angle pins and three dampers (D0, D1 and D2, Table 1), body pitch and aerodynamic damping of the flies were systematically varied. Results showed that for all individuals, forward velocity decreased sharply with body pitch (Fig. 2A), but only decreased slightly with damping coefficients 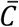 (Fig. 2B), except for the medium damping (D1) when *χ* = 22.5°. Note that the damping coefficients of medium (D1) and large (D2) damping cases were increased by 54% and 101% compared to that of small damping case (D0, no damper) (Table 1).

**Figure 2.**
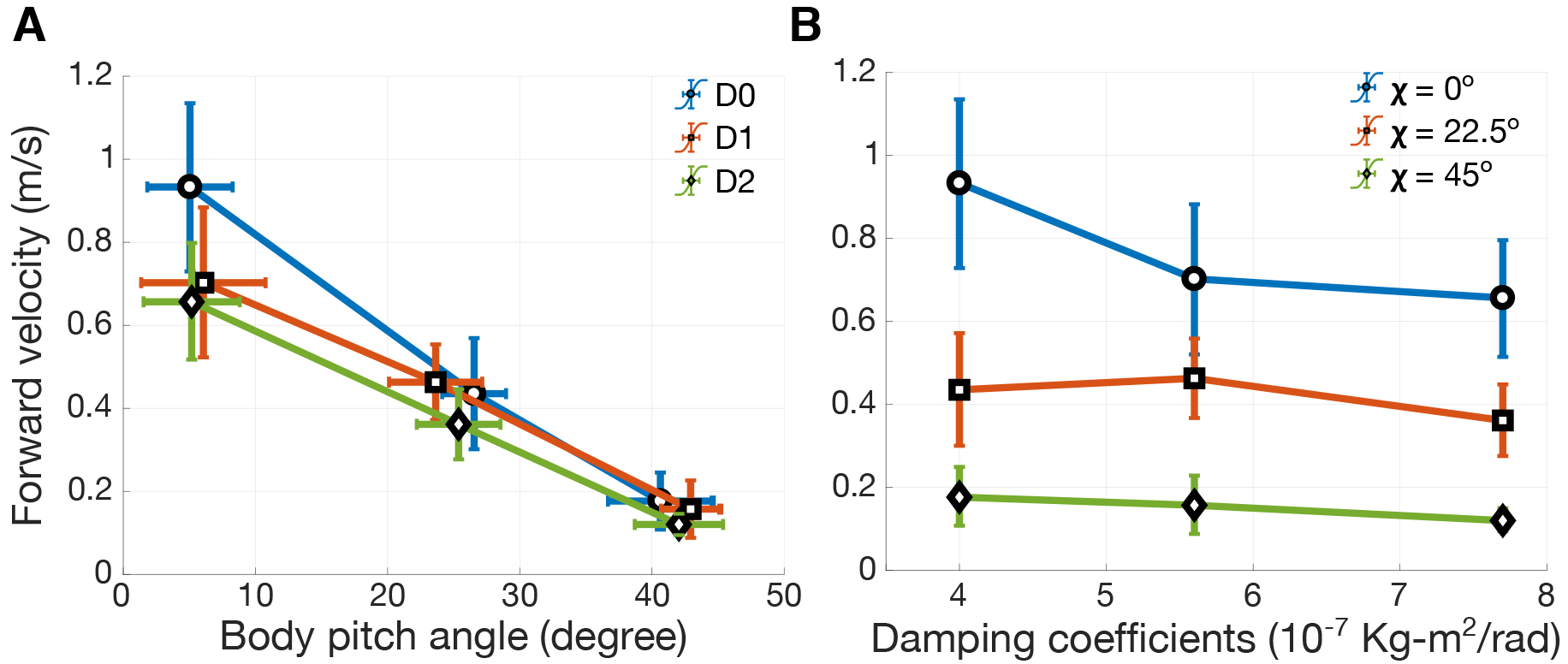
The dependency of forward velocity on body pitch angle and aerodynamic damping coefficient. (A) Forward velocity decreases approximately linearly with increasing body pitch angle in all three damping cases (D0 (blue), D1 (red), and D2 (green)). (B) Forward velocity decreases with increasing damping coefficients (except for *χ* = 22.5° and D1 damping case).

Although the dependency of forward velocity on body pitch angle and damping coefficients was consistent among individuals, there was also considerable variance of flight speed among individuals. For example, the slowest individual (BBF#1) cruised at a mean speed of 0.59 m/s in D0 case at *χ* = 0°, while the fastest individual (BBF#3) flew at 1.25 m/s under the same condition. It is also worth noting that all individuals performed smooth steady forward flight at lower body pitch angles (0° and 22.5°). However, at *χ* = 45° or above (not reported), the forward velocity reduced significantly to 0.15 ± 0.06 m/s and occasionally some flies produced vertical oscillations of the rotating shaft in the beginning of the trials. This was possibly due to the interaction between the wing lift force that tilted the MAGLEV pivot joint and the magnetic restoring torque due to the misalignment of the pivot permanent magnets and the parallel magnetic field generated by the electromagnets (Hsu et al., 2016). The oscillation usually diminished once steady-state flight has been reached.

### Wing kinematic variables during forward flight

As the body pitch, damping coefficient, and the resulting flight speed changed, there also existed considerable changes in wing kinematic patterns (Fig. 3 and Fig. 4). We characterized the changes using a collection of 9 wing kinematic variables representing the cycle-averaged features. The wing kinematic changes were more strongly correlated with the body pitch angle than with damping coefficients, as only wing deviation had noticeable correlation with the damping coefficients (Columns in Fig. 3). The contributions of these kinematic variables in speed control were tested according to the variable force-vectoring model (next section).

**Figure 3.**
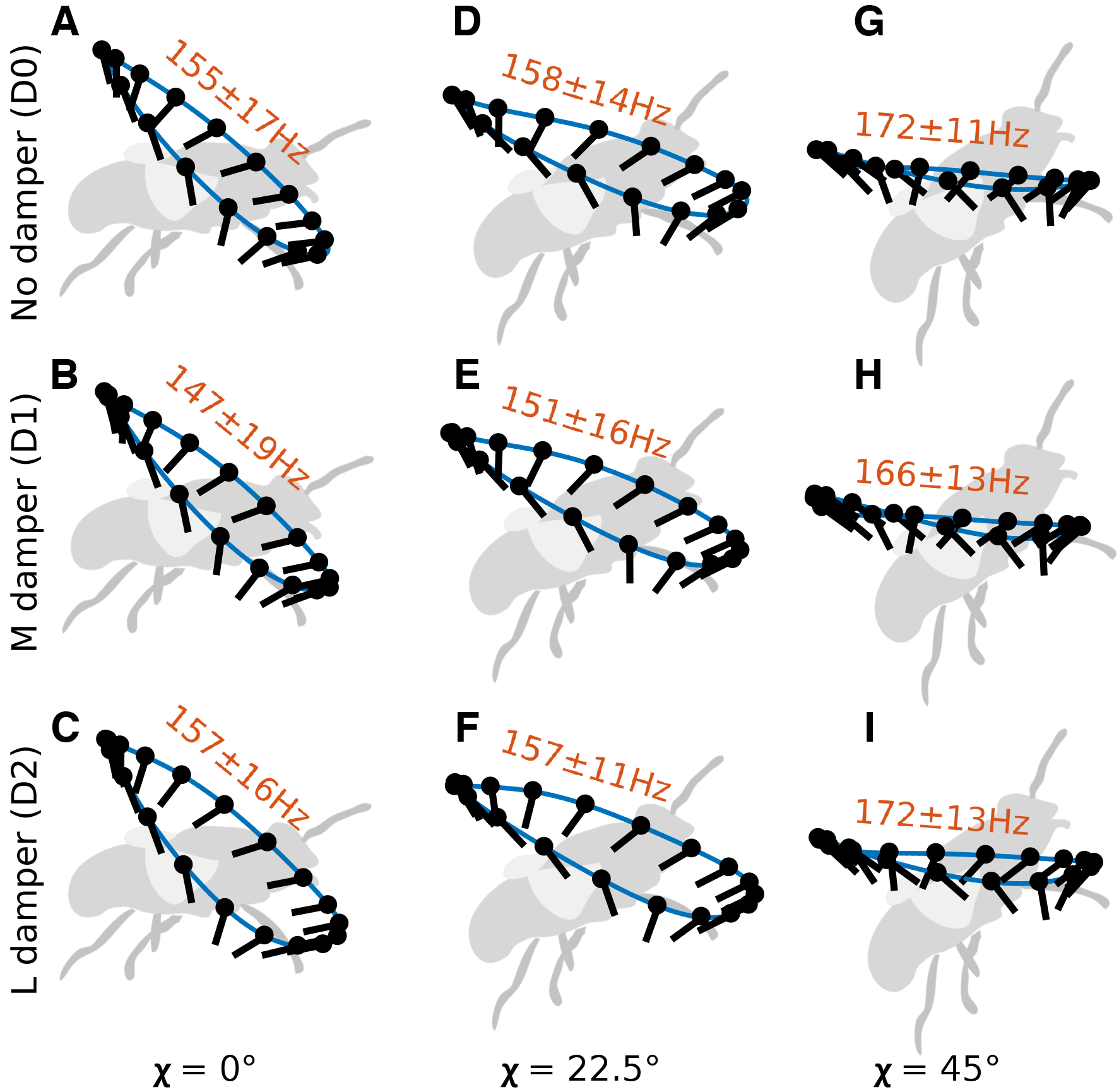
Averaged wingtip trajectories (blue curves) and mean wingbeat frequencies (The average and standard deviation are shown above each wingtip trajectory). At *χ* = 0° and *χ* = 22.5°, the wingtip trajectories are oval shapes, while at *χ* = 45°, the shape becomes flat. Stroke deviation amplitude (Φ) is the only variable that has noticeable correlation with both body pitch angle and damping coefficients (decreases with body pitch angle and increases with damper size).

**Figure 4.**
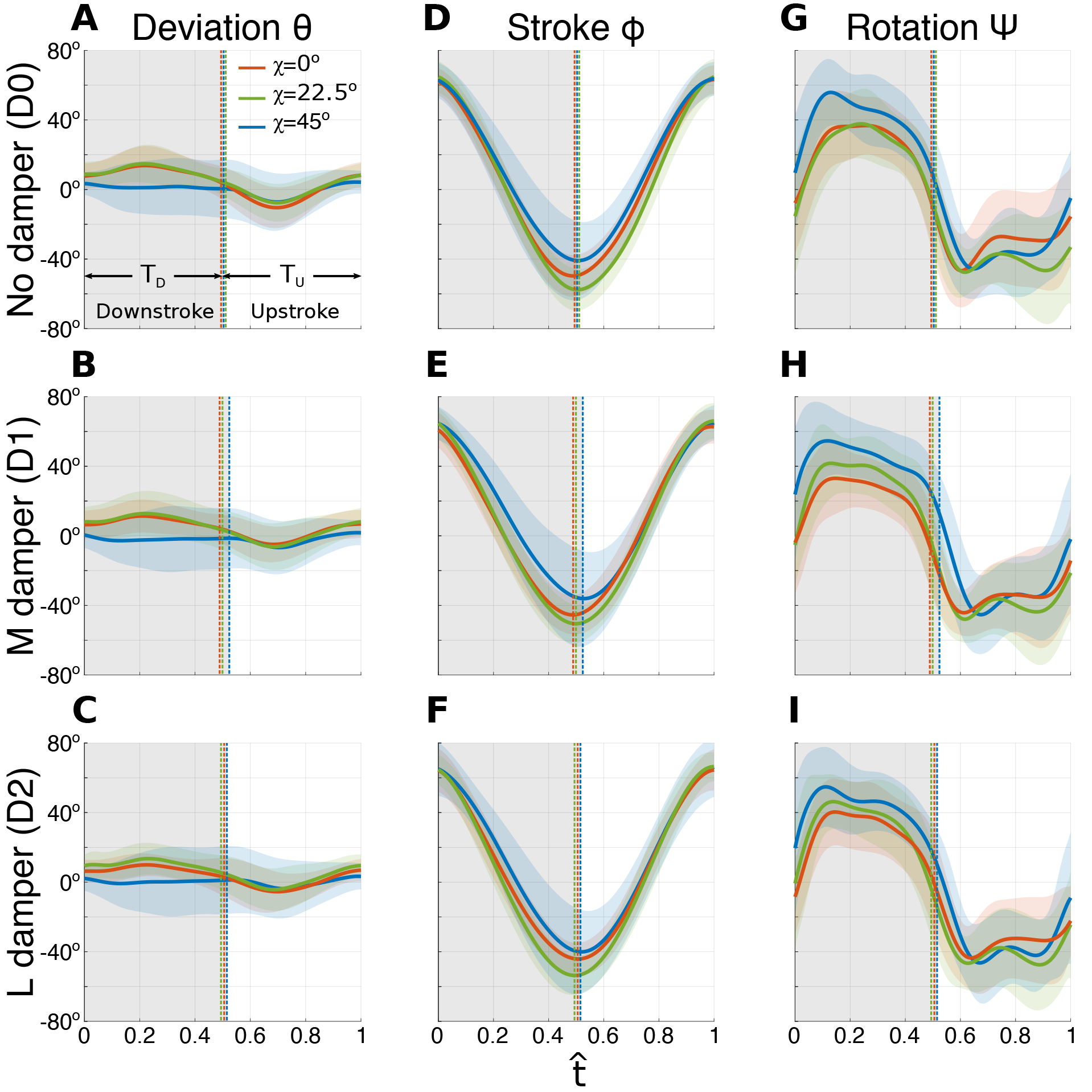
Traces of wing kinematic angles (deviation (*θ*), stroke (*ϕ*), and rotation (*ψ*)). The gray shades represent the duration of wing downstroke within one wingbeat 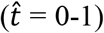. Colored (red: *χ* = 0°, green: *χ* = 22.5°, and blue: *χ* = 45°) shaded areas enclosing the curves indicate ±1 s.d. *T*_*D*_ and *T*_*U*_ are the downstroke and upstroke durations, respectively.

Mean wingbeat frequency (*n*) in each damping condition increased with body pitch angle (rows in Fig. 3), with approximately 11% increase from *χ* = 0° to *χ* = 45°. Therefore, wingbeat frequency had a clear decreasing trend with forward velocity. Mean wingbeat frequency averaged over all trials was 158.9 ± 16.6 Hz, with the highest at 172.4 ± 12.5 Hz (*χ* = 45° and D2 damping) and the lowest at 147.2 ± 18.7 Hz (*χ* = 0° and D0 damping). The ratio of downstroke and upstroke durations (T_*D*_/T_*y*_) generally increased with body pitch angle (Fig. 4A-C), as it peaked at 1.084 ± 0.077 at *χ* = 45° and D1 damping case and bottomed at 0.996 ± 0.065 at *χ* = 0° and D2 damping case.

Wing stroke amplitude (Φ), the changes of which mainly resulted from the extended excursion of the wing stroke towards the end of downstroke (forward excursion), was the largest at *χ* = 22.5° (123.3° ± 7.4°) and was the lowest at *χ* = 45° (111.1° ± 10.6°) (Fig. 4D-F). The mean wingtip velocity 2Φ*nR* (*R* as the wing length) dropped from 5.08 ± 0.59 m/s and 5.01 ± 0.62 m/s at *χ* = 22.5° and 45°, respectively, to 4.64 ± 0.76 m/s at *χ* = 0°.

Wing rotation amplitude (Ψ) increased with body pitch angle (except *χ* = 22.5° and D0 damping case). Maximum rotation angle (*ψ*_*max*_), occurred at the end of wing pronation, increased from 47.5° ± 10.4° at *χ* = 0° to 59.9° ± 12.8° at *χ* = 45° (Fig. 4G-I). Minimum rotation angle (*ψ*_*min*_), occurred shortly after the end of wing supination, bottomed at *χ* = 22.5° and rose approximately 5° for both *χ* = 0° and *χ* = 45° cases (Fig. 4G-I). Wing rotation angle at down-to-up stroke reversals (supination) tended to have a 10° - 14° delay relative to stroke at *χ* = 45°, while it was near symmetric (i.e., in phase with stroke) or slightly advanced at *χ* = 0° and 22.5° (except with D0 damping at *χ* = 0°) (Fig. 4G-I). Rotation angle at up-to-downstroke reversal (pronation) was advanced at *χ* = 45° and was delayed at *χ* = 0° and *χ* = 22.5°.

Wingtip trajectories at *χ* = 0° and *χ* = 22.5° took oval shapes and those at *χ* = 45° were more flat (Fig. 3 A-F for *χ* = 0° and *χ* = 22.5°; and G-I for *χ* = 45°). Deviation amplitude (Θ) decreased with increasing body pitch angle (Fig. 4A-C): 19.6° ± 8.9° at *χ* = 0° and 22.5°, and 12.8° ± 5.8° at *χ* = 45°. Deviation amplitude (Θ), which is the only kinematic variable that has noticeable correlation with damping coefficients, increased slightly with increasing damping coefficient, for example from 15.6° ± 4.7° with D0 to 24.0° ± 10.8° with D2 at *χ* = 0°. The increasing trend was less significant at *χ* = 22.5° and *χ* = 45° (Fig. 4A-C). A subtle decrease of wing stroke plane angle (*β* + *χ*) can be observed from *χ* = 0° to *χ* = 45° (rows in Fig. 3). The changes were limited, remaining within 44.5° ± 5° for all trials.

### Force-vectoring models for speed control and variable importance of wing kinematic variables

Two force-vectoring models for predicting the flight speed of flies were tested through nonlinear regression based on the estimated thrust, measured flight speed, and a collection of wing kinematic variables (described above). The nonlinear regression on constant force-vectoring model yielded total aerodynamic force magnitude *F*_0_ = 2.19 × 10^−4^ N with 95% confidence interval [1.99 × 10^−4^ N, 2.40 × 10^−4^ N] (or 53.2% [48.2%, 58.1%] of mean body weight) and *χ*_0_ = 47.8° [45.5°, 50.0°]. The RMSE of the prediction based on constant force-vectoring model was 0.131 m^2^/s^2^ with *R*^2^ of 0.71. It can be seen that the residual errors of constant force-vectoring model were relatively large as forward velocity increased (Fig. 5A).

**Figure 5.**
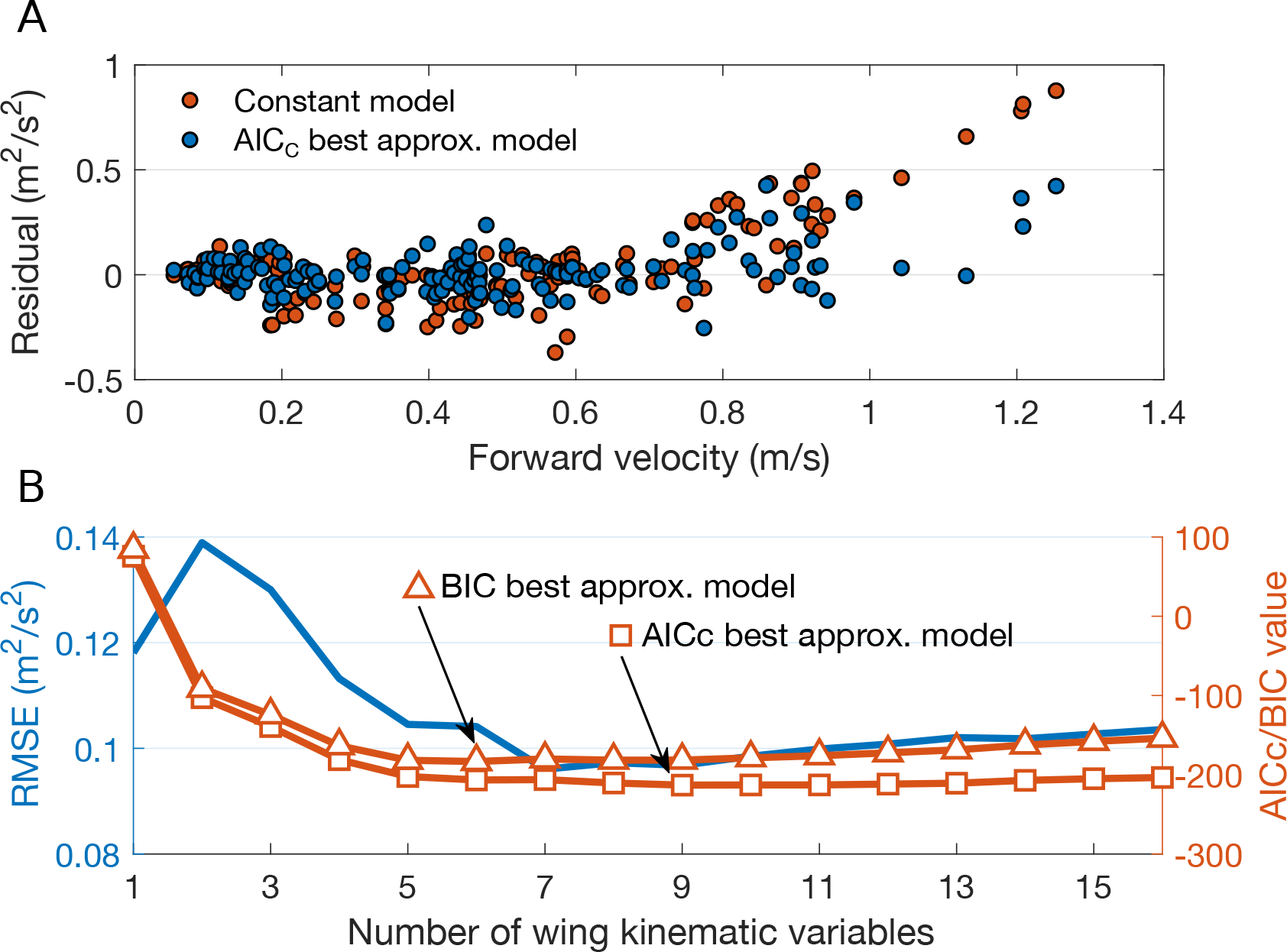
(A) Residual plot for the constant (red dots) and *AIC*_*c*_ best-approximating models (blue dots) with increasing forward velocity. (B) Graph of *AIC*_*c*_/*BIC* as a function of number of wing kinematic variables used in variable force-vectoring model. RMSE first goes up and quickly drops to minimum when the 7 of the most important variables are included (Ranking of the variable importance is shown in Fig. 6). *BIC* best-approximating model takes in 6 most important variables and *AIC*_*c*_ includes 9 variables.

**Figure 6.**
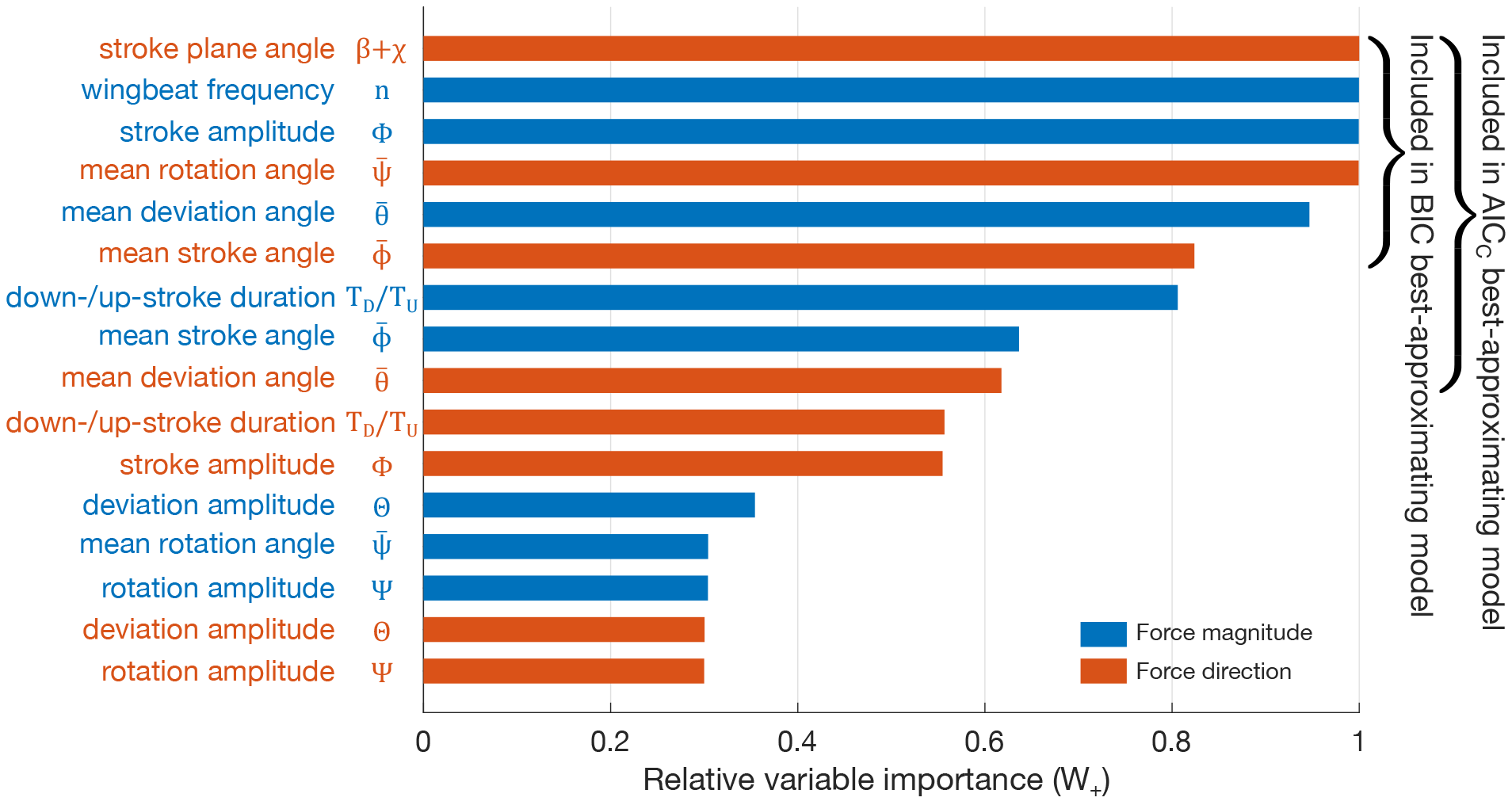
Variable importance of wing kinematic variables in the variable force-vectoring model (importance index based on summations of Akaike weights *w*_+_). Blue and red bars represent the relative importance of wing kinematic variables on force magnitude and direction, respectively. *BIC* and *AIC*_*c*_ best-approximating models include 6 and 9 most important variables, respectively.

The contributions of the 16 wing kinematic variables selected were tested in the variable force-vectoring model. Note that mean wingbeat frequency (*n*) and wing stroke plane angle (*β* + *χ*) were incorporated exclusively in force magnitude and force direction, respectively; and the other 7 wing kinematic variables, i.e., 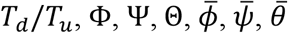 were included in both force magnitude and force direction. Nonlinear regressions were performed on all possible variable-combinations (total combinations: 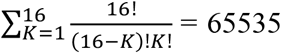. *AIC*_*c*_/*BIC* and Akaike weights (*w*) were computed for model selection and variable importance, respectively. From Fig. 5B, the *AIC*_*c*_ best-approximating model included 9 wing kinematic variables that contributed to the changes in the aerodynamic force magnitude and direction (Table 2); and *BIC* applied a larger penalty on model complexity, which reduced the variable number to 6 (Fig. 5B). *AIC*_*c*_ best-approximating model gave *F*_0_ = 3.04 × 10^−4^ N with 95% confidence intervals [2.85 × 10^−4^ N, 3.22 × 10^−4^ N] (or 73.7% [69.2%, 78.3%] of mean body weight), *χ*_0_ = 51.7° [50.6°, 52.9°], RMSE of 0.097 m^2^/s^2^, and *R*^2^ of 0.922. *BIC* best-approximating model gave *F*_0_ = 3.06 × 10^−4^ N with 95% confidence intervals [2.86 × 10^−4^ N, 3.25× 10^−4^ N] (or 74.2% [69.5%, 78.9%] of mean body weight), *χ*_0_ = 51.8° [50.7°, 52.9°], RMSE of 0.104 m^2^/s^2^, and *R*^2^ of 0.904. Both *AIC*_*c*_ and *BIC* best-approximating model reduced the residual error compared to constant force-vectoring model, particularly in higher velocity range (Fig. 5A). Here, we chose *AIC*_*c*_ model as our best-approximating model which biased slightly towards the goodness-of-fit than the *BIC* and included 3 additional wing kinematic variables (Fig. 6).

**Table 2.**
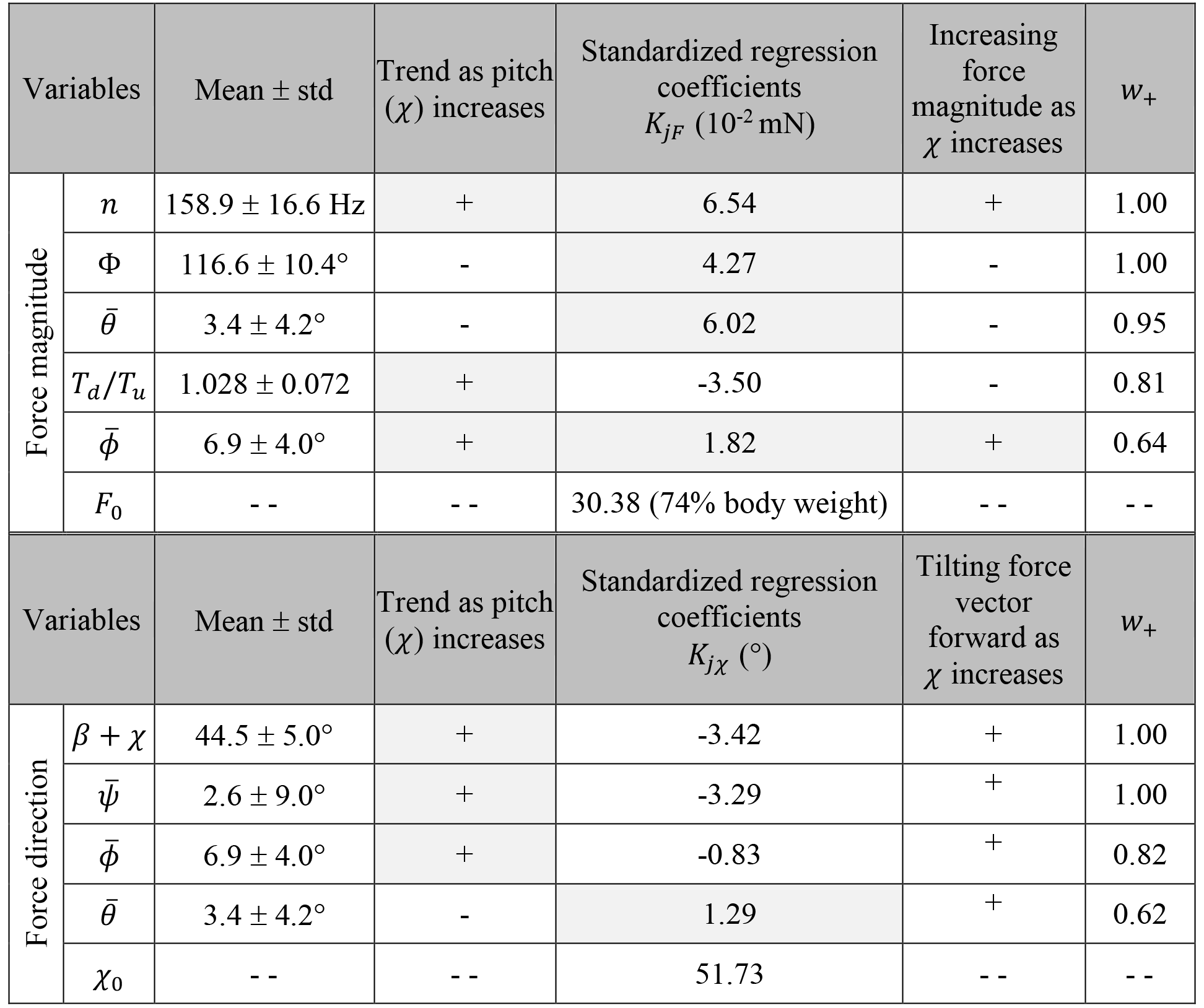
Wing kinematic variables that modulate force magnitude and direction as body pitch angle changes. The table summarizes their mean values, standard derivation, trend (+ increasing, − decreasing) as body pitch increases, whether they increase force magnitude or tilt force vector forward as body pitch increases, and standardized regression coefficients *K*_*jF*_ and Akaike weight *w*_+_ from the nonlinear regression result of *AIC*_*c*_ best-approximating model. The wing kinematic variables that contribute to force magnitude (Δ*F*) in Eqn. 9 are: mean wingbeat frequency (*n*), stroke amplitude (Φ), mean deviation angle 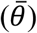, ratio of downstroke and upstroke durations (*T*_D_/*T*_*U*_), and mean stroke angle 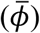; and *F_0_* is the constant term in force magnitude. The wing kinematic variables that contribute to force magnitude (Δ*χ*) in Eqn. 9 are: stroke plane angle (*β* + *χ*), mean rotation angle 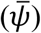, mean stroke angle 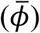 and mean deviation angle 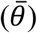; and *χ*_0_ is the constant term in force angle. The trend of a variable as pitch *χ* increases is calculated based on the Pearson’s bivariate correlation and is marked as + (increasing) or − (decreasing). *w*_+_ is the summation of Akaike weights of each wing kinematic variable. A positive sign of *K*_jF_ indicates that an increase of the kinematic variable directly increases the force magnitude or tilts the force backward (independent of *χ*).

Next, with the summations of Akaike weights (*w*_+_), the relative importance for the wing kinematic variables is shown in Fig. 6. Wing kinematic variables that contributed to the force magnitude were: 1) wingbeat frequency (*n*), 2) stroke amplitude (Φ), 3) mean deviation angle 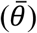, 4) ratio of downstroke and upstroke durations (*T*_*D*_/*T*_*u*_), and 5) mean stroke angle 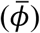. Among these variables, force magnitude Δ*F* (in Eqn. 9) depended negatively on *T*_*D*_/*T*_*u*_, meaning smaller duration of downstroke period increases the Δ*F*, while higher wingbeat frequency and amplitude, upward shift of mean deviation angle (so that wing trajectory becomes more oval) and dorsal (backward) shift of mean stroke angle all led to higher Δ*F* (see signs of *K*_*jF*_ in Table 2). Kinematic variables that contributed to force direction were: 1) stroke plane angle (*β* + *χ*), 2) mean rotation angle 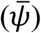, 3) mean stroke angle (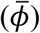), and 4) mean deviation angle 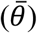. Among these variables, force direction (Δ*χ* in Eqn. 9) depended positively only on 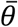, meaning upward shift of mean deviation angle (or more oval wing trajectory) results in a backward tilt of force direction (Δ*χ* increases), while the increases in *β* + *χ* (i.e., forward tilt of stroke plane), 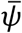 (i.e., increased pronation/decreased supination), and 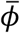 (i.e., backward shift of wing stroke angle), all result in a forward tilt of the force direction (Δ*χ* decreases) (see signs of *K*_*jχ*_, Table 2).

As the body pitch angle *χ* increased and flight speed decreased, changes in *n* (increase) and 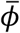 (increase) led to increases on force magnitude (see effect on force magnitude as *χ* increases, in Table 2), while the changes in Φ (decrease), 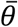 (decrease), and *T*_*d*_/*T*_*u*_ (increase), led to the decrease in force magnitude (Table 2). In total, the force magnitude increased slightly with the increasing pitch angle, as the collective result of all wing kinematic changes. In addition, as the body pitch angle increased, all the changes of wing kinematic variable resulted in a forward tilt of force direction (see effect on force direction as *χ* increases Table 2) (*β* + *χ* increases, 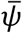 increases, 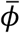 increases and 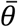 decreases), thereby to compensate the thrust loss due to the backward force tilt.

## DICUSSION

### Flies control forward velocity using their equivalent to the helicopter control

Not surprisingly, the inverse dependency between the forward velocity (*ν*) and the body pitch angle (*χ*) (Fig. 2A) in blue bottle flies is consistent with those observed in other insect species (Azuma and Watanabe, 1988; David, 1978; Dudley and Ellington, 1990; Meng and Sun, 2016; Willmott and Ellington, 1997) and birds (Brown, 1963; Pennycuick, 1968). This suggests that blue bottle flies mainly rely on body pitch adjustment to vector the wing aerodynamic forces to produce thrust and regulate flight speed. However, the current study reveals more intricacies in the force-vectoring and speed control of flies, which show close resemblance to those of helicopters, or to the “helicopter model”. Helicopters create thrust and pitch moment using cyclic and collective pitch, in conjunction with throttle (Leishman, 2006). Collective pitch and throttle increase the force magnitude by symmetrically increasing the blade AoA and engine speed, respectively. Cyclic pitch tilts the rotor disc and aerodynamic force forward through precession effect and blade flapping caused by asymmetric modulation of blade AoA (Leishman, 2006). This produces a forward thrust and a pitch moment that tilts the helicopter body forward. In this process, the tilt of the rotor disc is relatively small and less conspicuous than the tilt of the helicopter body itself; as a result, the angle between the aerodynamic force and the helicopter body is only modulated within a limited range, and the total vectoring of the aerodynamic force is determined mainly by the body pitch. This gives rises to the so-called helicopter model, where the thrust and speed are mainly determined by body pitch and throttle.

Through testing constant and variable force-vectoring models, here we show that blue bottle flies flying in the flight mill closely follow the helicopter model. First, the constant force-vectoring model (Eqn. 7), assuming the thrust and forward velocity are determined solely by body pitch, yields a reasonable prediction of the forward velocity (R^2^ = 0.71, RMSE = 0.131 m^2^/s^2^), confirming the dominant role of body pitch in speed control. Next, the variable force-vectoring model (Eqn. 9) further reveals the importance of wing kinematic control in predicting the flight speed, similar to the role of collective pitch, cyclic pitch, and throttle of helicopters. Specifically, the magnitude of the aerodynamic force is controlled by mean deviation angle, ratio of downstroke and upstroke duration, and mean stroke angle, which can be seen as flies’ equivalent of collective pitch (Fig. 6). In addition, stroke amplitude and wingbeat frequency also control the force magnitude, which resemble the function of throttle, or the “engine speed” of the helicopters. The force direction, on the other hand, is controlled primarily by stroke plane angle, mean rotation angle, mean stroke angle, and mean deviation angle (Fig. 6), which can be seen as flies’ equivalent of cyclic pitch. The results also show that these kinematic variables collectively lead to moderate modulation of force magnitude (Δ*F* = 8.5 × 10^−5^ ± 7.2 × 10^−5^ N, or 20.5 ± 17.5% of mean body weight in Eqn.9), but only minor change in force vector direction (Δ*χ* = 3.76± 2.77° in Eqn. 9), which resembles closely to the speed control of helicopter.

In summary, our results show that although flies use flapping wings instead of rotary wings, and are capable of large modulation of wing kinematic (Deora et al., 2015), their thrust generation mechanism and flight speed control still conform to the helicopter model with limited change of wing kinematics, at least within the range of speeds achieved while flying in the MAGLEV flight mill. Large modulation of wing kinematics presumably only occur during short-period transient flight such as landing on the ceiling (unpublished data from authors) and recovery from extreme perturbations (Beatus et al., 2015). Finally, note that the contribution of body pitch on speed control may subject to saturation at higher speed, where wing kinematic modulation becomes the primary mechanism. For example, a recent study (Meng and Sun, 2016) show that drone-fly can fly at a wide range of speed (3.1m/s to 8.4 m/s) for a brief amount of time prior to landing at almost the same body pitch (close to zero degrees), and relatively large changes in wing kinematics are employed by the flies to regulate speed.

### Physical significance of wing kinematic variables identified in the *AIC*_*c*_ best approximating model

In variable force-vectoring model, we have identified a collection of wing kinematic variables (Fig. 6), which represent either symmetric or asymmetric changes of wing motion between half-strokes. In our experiments, these kinematic variables changed in response to the changes of body pitch angle, together they control the flight speed of the flies through altering the aerodynamic force magnitude and direction. The specific roles of each wing kinematic variables in modulating the force magnitude and direction are quantified by the regression coefficients *X*_*jF*_ and *X*_*jχ*_ in Table 2. The trend of their changes in response to body pitch angle can be quantified through Pearson’s bivariate correlation between each wing kinematic variables and the body pitch, and the signs of the regression coefficients are summarized in Table 2, together with the magnitude of their changes quantified by their standard derivation. With these results, here we discuss the physical significance of each wing kinematic variables in force modulation and speed control.

The magnitude of the aerodynamic force is mainly controlled by wingbeat frequency, stroke amplitude, mean deviation and stroke angles, and the ratio of downstroke/upstroke duration (Fig. 6 and Table 2). Wingbeat frequency increased by 11% on average from *χ* = 0° to *χ* = 45° (rows in Fig. 3), indicating that flies were attempting to increase force magnitude and therefore to compensate thrust loss while the force vector is being tilted backward with increasing pitch angle. This relatively small increase of wingbeat frequency is expected for flies with asynchronous power muscles, as the wingbeat frequency is primarily determined by the mechanical properties of the coupled wing-and-thoracic oscillator, which only permits slight alteration of wingbeat frequency (Bartussek et al., 2013). In addition, flies decrease the stroke amplitude Φ as *χ* increases, which is accompanied by a backward shift of mean stroke angle 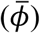, through reducing the forward excursion (Fig. 4D-F). Since the increases of the increases of both variables increase force magnitude, the decreasing trend of stroke amplitude and increasing trend of mean stroke angle, result in opposite effects on force magnitude when body pitch increases (Table 2). The backward shift of mean stroke angle also tilts the force vector forward (Table 2) and creates a pitch down torque at high pitch angle (indicating the flies are attempting to lower its body pitch to compensate thrust loss). Flies also increased the duration of upstroke (wings sweep backward), during which the thrust is mainly generated, and reduced the duration of downstroke (wings sweep forward), during which drag is mainly generated; together they both increase the total force magnitude at lower body pitch.

The change in mean deviation angle 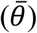 is also a strong contributor to force magnitude, while also being a contributor to force direction. At higher flight speed (or lower pitch), there is an increase of 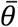, mainly results from the increase of deviation during downstroke (Fig. 4A-C), which renders the shape of the wingtip trajectory more oval. The oval shape introduces a velocity component perpendicular to the mean stroke plane, upward during downstroke and downward during upstroke. As suggested by Sane and Dickinson (2001), upward velocity reduces AoA and drag force during downstroke and downward velocity results in an increase in AoA and thrust during upstroke, together they increase force magnitude.

The direction of the aerodynamic force is mainly controlled by the stroke plane angle, and mean stroke, rotation and deviation angles (Fig. 6 and Table 2). At higher body pitch angles, blue bottle flies increase (*β* + *χ*) and decrease 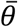 (rows in Fig. 3), although in small variations (44.5° ± 5.0° and 3.4° ± 4.2°), to tilt the stroke plane and the aerodynamic forces more forward as body pitch angle increases, which are clear signs of attempting to compensate the loss of thrust. At large pitch angles, they also increase the mean rotation angle (shifted forward from 8° to 16°), this increases and decreases the AoA during upstroke and downstroke respectively, and also tilt the force vector forward. In summary, we find all 4 wing kinematic variables contributed to the force direction tend to compensate the loss of thrust as the body pitches up.

### Left and right wing asymmetry

In the analysis of wing kinematic variables responsible for forward flight, we averaged left and right wing kinematics. However, asymmetry was observed between left and right wing motion due to the rotational nature of the flight mill. For example, mean right (inner) wing stroke amplitude (Φ_*r*_) was 13° higher than mean left (outer) wing stroke amplitude (Φ_*l*_). A likely explanation is that the halteres - organs modified from hindwings, unique to *Diptera*, measure the angular rate (Dickinson, 1999; Taylor and Krapp, 2007) - sensed the difference between left and right wings and tried to initiate a body yaw turn (Dickinson, 1999). Another possibility for the wing asymmetry could be due to the blue bottle flies’ tendency to perform corrective yaw turns to balance the optical flow experienced by the left and right compound eyes (29 out of 154 total trials, Φ_*l*_ > Φ_*r*_). Note that, due to the use of identical spatial frequency of the grating patterns on the two walls, the outer wall had higher temporal frequency because of the larger radius. It has been shown that honeybees use a “centering response” to mediate the unbalanced optical flow (Srinivasan and Zhang, 2004). Subtle stroke amplitude difference between left and right wing can produce roll and yaw moment (Fry et al., 2003), the 13° difference observed here is higher than those observed in free flight saccades in fruit flies, which is likely to result from either continuous yawing or saturated saccadic responses (because of the tether, the error in yaw or roll control cannot be compensated, and any integral controller tends to saturate the response (Muijres et al., 2015)). Nevertheless, the existence of the asymmetry does not prevent us from analyzing the forward flight by averaging the left and right wing kinematics, and interestingly it also shows the potential of using the flight mill to investigate a fly’s speed control and yaw responses to (bilaterally asymmetric) visual stimuli together with mechanosensory inputs in steady forward flight.

### The advantages and limitations of experiments using MAGLEV flight mill

In this study, we demonstrated a novel design and use of MAGLEV flight mill in studying voluntary steady forward flight of blue bottle flies. Instead of “forcing” the flies to fly under a prescribed freestream airflow in wind tunnels, the blue bottle flies flew voluntarily with a speed resulted from its own effort and sensorimotor response. Further, the magnetically-levitated pivot joint eliminated the mechanical friction compared to traditional flight mills with mechanical pivots, therefore allowing easier and more accurate calibration and manipulation of aerodynamic damping that the flies had to overcome. Together with magnetic angle pins to vary body pitch angle and the enclosed cylinder walls covered by grating patterns to keep consistent visual stimuli, this MAGLEV flight mill enabled semi-free flight experiments in a well-controlled environment. Therefore, this device could be further exploited to investigate insects’ aerodynamics, dynamics, sensing and control in forward flight.

Compared with the free flight experiments using wind tunnels, there are also a few limitations of the MAGLEV flight mill. First, due to the existence of aerodynamic damping acting on the rotating rod, which cannot be fully eliminated, the fastest steady-flight speed of the blue bottle flies observed in the experiments was 1.25 m/s, which is slower than that in free flight. However, flies in free flight hardly reach a steady-state. For example, flying in a 1.6 m^3^ chamber, blue bottle flies showed top speed at 2.5 m/s and an average speed at 1.3 m/s with constantly accelerate and decelerate (Bomphrey et al., 2009). Second, studies showed that tethering may significantly reduce wingbeat frequency (Baker et al., 1981; Betts and Wootton, 1988; Kutsch and Stevenson, 1981). However, no significant difference in wingbeat frequency was observed between the current study (158.9 Hz in average) and typical blowflies (150Hz) (Dickinson, 1990). Likewise, the same conclusion was made in beetles’ forward flight in another flight mill study (Ribak et al., 2017). Third, the rotational nature of the flight mill has caused noticeable bilateral wing asymmetries as described above. While here we considered the effect of these asymmetries negligible on the forward speed, they need to be limited to certain degree by using sufficiently large rod radius. Nevertheless, as described above, they can be possibly exploited to study the role of visual and mechanosensory feedback in forward flight. Lastly, lift was significantly reduced (mean lift is 69.5 % of mean body weight) since flies did not have to actively maintain aloft. This reduction could be slightly remedied by banking the roll axis during circular flight to balance the centrifugal force (Ribak et al., 2017). Nonetheless, holding the effects of these limitations in check, this device provides an alternative approach to a wind tunnel to study insect forward flight in controlled conditions and has large potential to be further exploited in the future as a common tool in insect flight research.

## LIST OF SYMBOLS AND ABBREVIATIONS

*A*: area
*AIC*: Akaike information criterion
*AIC*_*c*_: small sample AIC
*BIC*: Bayesian information criterion
*β*: stroke plane angle relative to horizontal
*β*+*χ*: stroke plane angle relative to body long axis
*C*_*D*_: drag coefficient
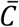: average damping coefficient
D1~D3: no damper, medium damper, and large damper
Δ_*i*_: AIC difference of *i*^*th*^ model
*F*: Resultant force
*F*_*thrust*_: thrust force
*F*_lift_: lift force
γ: angle pin angle
*I*: moment of inertia
*k*: constant of integration
*K*_jF_, *K*_j*χ*_: standardized regression coefficients of wing kinematic variables contribute to force magnitude and force direction
*K*: number of parameter in the candidate model
*l*: radius of flight mill
*m*: total number of trials
*n*: wingbeat frequency
*Φ*: stroke amplitude
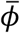: mean stroke angle
*ϕ*_0_, *ϕ*_*si*_, *ϕ*_*ci*_: constant, sine, and cosine Fourier coefficients of stroke position
Ψ: rotation amplitude
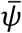: mean rotation angle
*ψ*_0_, *ψ*_*si*_, *ψ*_*ci*_: constant, sine, and cosine Fourier coefficients of wing rotation
*ρ*: mass density of the fluid
*R*: wing length
*R*^2^: coefficient of determination
*S*: first moment of area
*t*: time
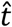: dimensionless time
T_*D*_/T_*U*_: ratio of downstroke and upstroke durations
Θ: deviation amplitude
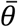: mean deviation angle
*θ*_0_, *θ*_*si*_, *θ*_*ci*_: constant, sine, and cosine Fourier coefficients of stroke deviation
τ: torque
*ν*: body forward velocity
*ω*: flight mill angular velocity
*w*: Akaike weight
*w*_+_: summation of Akaike weights
*χ*: body pitch angle
*χ*_0_: angle between total force vector and body pitch axis
*X*_*jF*_, *X*_*jχ*_: wing kinematic variables contribute to force magnitude and force direction
(*X, Y, Z*): global coordinate frame
(*X*_*b*_, *Y*_*b*_, *Z*_*b*_): body coordinate frame
(*X*_*S*_, *Y*_*S*_, *Z*_*S*_): stroke plane coordinate frame
(*X*_*w*_, *Y*_*w*_, *Z*_*w*_): wing coordinate frame

## AUTHOR CONTRIBUTIONS

S.J.H. and B.C. contributed to the design of the experiments. S.J.H and N.T. prepared and conducted the experiments and contributed to the data analyses. All authors contributed to the interpretation of the findings and the manuscript preparation.

